# A Human 3D neural assembloid model for SARS-CoV-2 infection

**DOI:** 10.1101/2021.02.09.430349

**Authors:** Lu Wang, David Sievert, Alex E. Clark, Hannah Federman, Benjamin D. Gastfriend, Eric Shusta, Sean P. Palecek, Aaron F. Carlin, Joseph Gleeson

**Affiliations:** Department of Neurosciences, University of California San Diego, La Jolla, CA, 92093, USA; Rady Children’s Institute for Genomic Medicine, Rady Children’s Hospital, San Diego, CA, 92123, USA; Department of Medicine; University of California San Diego, School of Medicine, La Jolla, CA, 92093, USA; Center for Immunity and Inflammation, New Jersey Medical School, Rutgers-The State University, Newark, NJ, 07103, USA; Department of Chemical and Biological Engineering, University of Wisconsin-Madison, Madison, WI, 53706, USA; Department for Pediatrics, University of California San Diego, La Jolla, CA, 92093, USA

**Keywords:** Cortical brain Organoid (CO), PLCs Containing Cortical Organoid (PCCO), SARS-CoV-2, Astrocytes, Viral replication hub

## Abstract

Clinical evidence suggests the central nervous system (CNS) is frequently impacted by SARS-CoV-2 infection, either directly or indirectly, although mechanisms remain unclear. Pericytes are perivascular cells within the brain that are proposed as SARS-CoV-2 infection points^1^. Here we show that pericyte-like cells (PLCs), when integrated into a cortical organoid, are capable of infection with authentic SARS-CoV-2. Prior to infection, PLCs elicited astrocytic maturation and production of basement membrane components, features attributed to pericyte functions in vivo. While traditional cortical organoids showed little evidence of infection, PLCs within cortical organoids served as viral ‘replication hubs’, with virus spreading to astrocytes and mediating inflammatory type I interferon transcriptional responses. Therefore, PLC-containing cortical organoids (PCCOs) represent a new ‘assembloid’ model^2^ that supports SARS-CoV-2 entry and replication in neural tissue, and PCCOs serve as an experimental model for neural infection.

## Main

Initially thought of as primarily a respiratory infection, SARS-CoV-2 is now implicated in substantial central nervous system (CNS) pathology^3,4^. CNS symptoms include ischemic strokes, hemorrhages, seizures, encephalopathy, encephalitis-meningitis, anosmia, post-infectious syndromes, and neuro-vasculopathy, collectively described in up to 85% of ICU patients^5–8^. Several reports appear to meet established criteria for infectious encephalitis^9^.

SARS-CoV-2 can utilize angiotensin-converting enzyme 2 (ACE2) as a receptor although other receptors have been proposed^10,11^. Recent studies on single cell RNA-seq (sc-RNA-seq) datasets indicate low levels of ACE2 expression in brain cells, however, expression is relatively high in some neurovascular unit (NVU) components, particularly in brain pericytes^12–14^. Autopsy series have suggested the potential for SARS-CoV-2 to spread throughout the brain, especially within vascular and immune cells. They note ischaemic brain lesions accompanied by widespread activation of astrocytes and cell death^1,15^. The potential for a SARS-CoV-2 elicited neurovasculopathy supports the development of new models to study tropism and pathology.

Brain pericytes are derived from neural crest stem cells (NCSCs), and are uniquely positioned in the neurovascular unit (NVU), physically linking endothelial and astrocytic cells^16^. Embedded within the basement membrane, pericytes connect, coordinate and regulate signals from neighboring cells to generate responses critical for CNS function in both healthy and disease states, including blood-brain barrier permeability, neuroinflammation, neuronal differentiation, and neurogenesis in the adult brain^17–19^.

We found that GFP^+^ pericyte-like cells (PLCs) generated *in vitro* from human pluripotent stem cell (hPSC)-derived NCSCs expressed the standard pericyte markers NG2 and PDGFR-β (Fig. 1a-b)^20^. We detected appreciable ACE2 mRNA and protein in 2D cultured PLCs compared with cultured human neural precursors (Extended Data Fig. 1a-d). To assess SARS-CoV-2 PLC tropism, we exposed PLCs to authentic SARS-CoV-2 at MOI 0.5, then harvested supernatant and cells daily (Fig. 1c). We found that the percentage of SARS-CoV-2 nucleocapsid (SNP)-positive cells and viral RNA as measured by qRT-PCR increased daily up to 72 hours post infection (h.p.i.), from 0% to 65% SNP positive, with viral RNA load increasing up to ~1000-fold (Fig. 1d-e). Plaque assay from supernatants on Vero E6 cells showed ~100-fold increased infectious virus production at 24 h.p.i with increased virus particles as well as viral titers compared to baseline, suggesting viral production by PLCs (Fig. 1f-g)^21^.

**Fig. 1.**
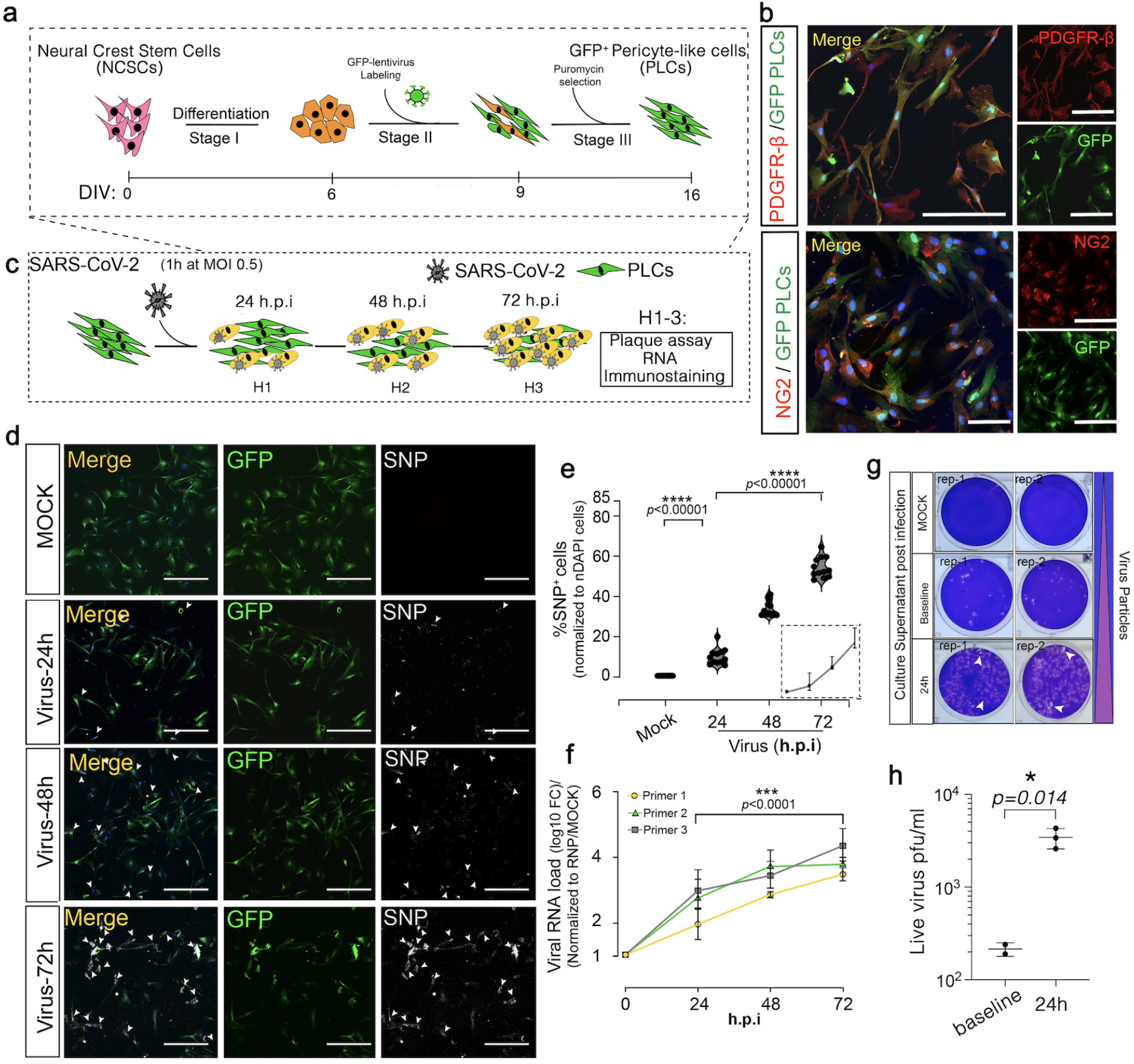
SARS-CoV-2 productively infects pericyte-like cells. **a.** GFP^+^ pericyte-like cells (PLCs) generation protocol. **b.** Immunostaining of GFP^+^ PLCs shows expression of pericyte markers PDGFR-β, and NG2. GFP^+^ PLCs cultured on coverslips were fixed and immunostained. Blue: DAPI, bar: 100μm. **c.** SARS-CoV-2 infection of PLCs protocol. At each time-point, RNA was isolated and in parallel cells were fixed and immunostained. **d.** SARS-CoV-2-Nuclecapsid protein (SNP) co-visualized with GFP. DAPI: nuclei, Bar: 400μm. GFP: PLCs, Arrows: SNP^+^ cells. **e.** Quantification of SNP expression from d. SNP^+^ cells were counted in ImageJ, nDAPI used as reference. n=12 includes 3 PLC cover slips from different biological replicates and 4 image regions from each cover slip. t-test with Sidak multiple-comparison correction was used to determine the significance. **** p<0.00001. h.p.i: hours post infection. **f.** qRT-PCR for SARS-CoV-2 shows increasing viral RNA load over time. *RNP* used as reference. n=12 includes PLCs from 3 different wells and 4 technical replicates for each biological replicate. t-test with Sidak multiple-comparison test correction was used to calculate significance *** p<0.0001. h.p.i: hours post infection. **g.** Plaque assay shows increased viral particles in culture supernatant 24h post infection. MOCK and baseline infection used as negative control. Two replicates (rep-1, rep-2) from each condition are shown. **h.** Plaque assay indicates secretion of live virus into culture supernatant at 24h post infection; n=3 includes three replicates generated from three different biological replicates for 24h, and n=2 for baseline infection, p value = 0.014, * p<0.1. Two-way ANOVA followed by a Sidak multiple-comparison test was used for statistics.

Demonstrating the infectability of PLCs led us to explore their effects on SARS-CoV-2 tropism within cortical brain organoids (COs). To this end, we developed an ‘assembloid’ in which GFP^+^ PLCs are integrated into mature COs. We thus generated PLC-containing cortical organoids (PCCOs), by seeding 2×10^5^ GFP^+^ PLCs into wells containing COs at 60div (days in vitro) (Fig. 2a). By 74div, using tissue clearing and light sheet microscopy, we observed GFP^+^ cells integrating into COs as cell clusters, subsequently spreading across the surface and penetrating into the CTIP2^+^ cortical plate-like zone in COs (Fig. 2b-c).

**Fig. 2.**
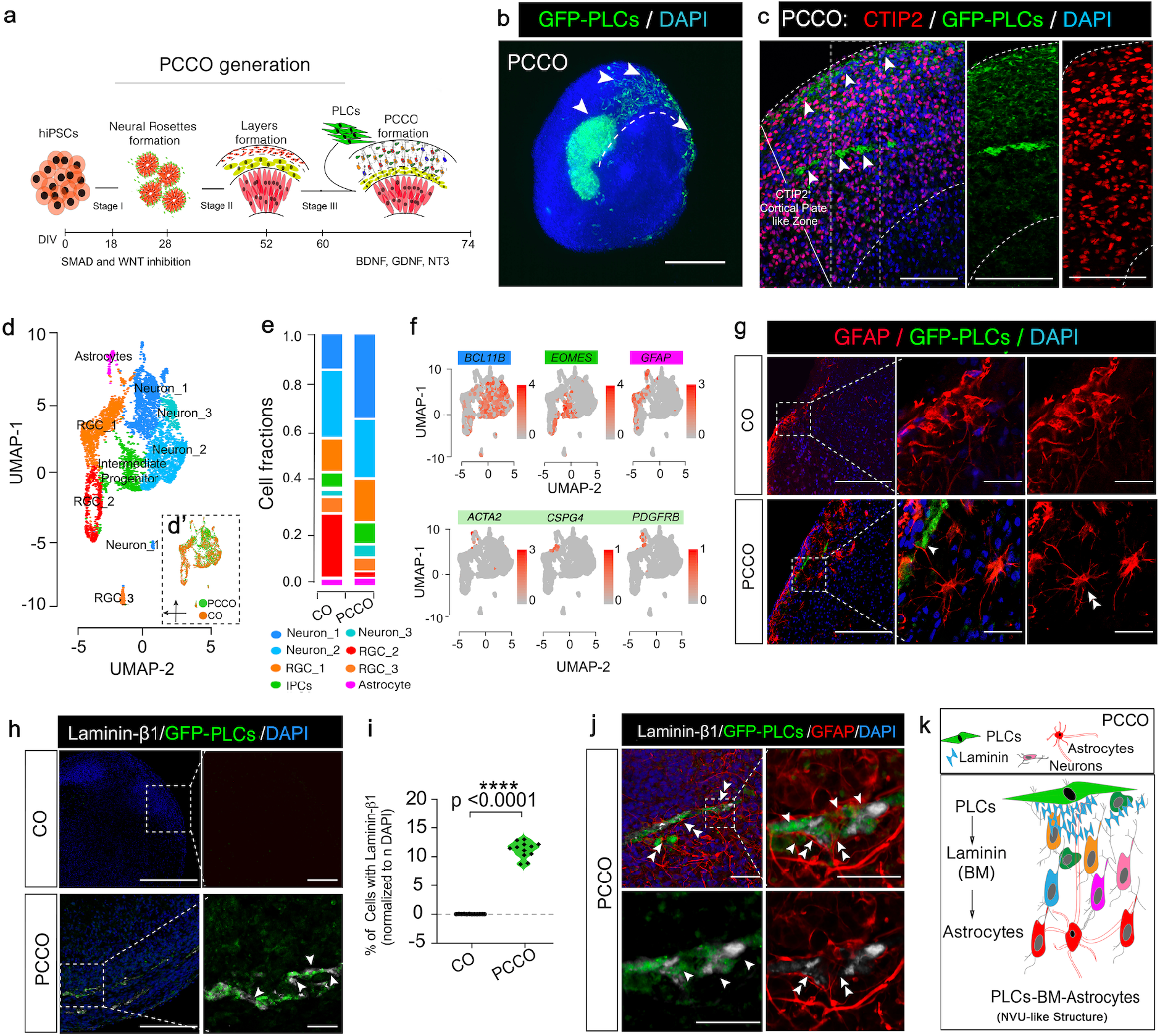
PCCOs incorporate pericyte-like cells into cortical organoids. **a.** PCCO generation protocol. **b.** Light-sheet image of GFP^+^ pericyte-like cells (PLCs) invading cortical organoid. Bar: 800μm, arrow: PLCs. **c.** GFP^+^ PLCs penetration into the CTIP2^+^ cortical plate (CP) like zone. DAPI: nuclei. Bars: 400μm, arrows: PLCs. **d.** Merged UMAP shows PCCO cellular compositions; 8 cell clusters annotated according to standard marker gene expression. d’: split UMAP shows the contribution of CO and PCCO. **e.** Cell fraction plot for proportionate cell clusters in CO and PCCO. IPC: intermediate progenitor cells; Astro: astrocytes. **f.** Features plots for CO and PCCO. *BCL11B:* deeper layer neurons, *EOMES:* intermediate progenitors, *GFAP:* astrocytes, *ATCA2, CSPG4(NG2), PDGFRB:* PLCs. **g.** ‘Star’-shaped astrocytic morphology seen in PCCOs but not COs. GFP: PLCs, bar: 400 μm, zoom-in bar: 100μm, double arrow: astrocytic endfeet connection, arrow: star-shaped astrocytes. **h.** Laminin-β1 expression and co-localization adjacent to GFP^+^ PLCs, evident only in PCCOs. Bar: 400μm, zoomin bar: 100μm. Single white arrow: laminin in gray. **i.** Quantification for H. Cells with adjacent laminin-β1 were counted in ImageJ. DAPI+ (nDAPI) were used as reference; n=12 includes 3 PCCOs from independent biological replicates and 4 image regions from each section. t-test with Sidak multiple-comparison test correction was used to determine the significance **** p<0.0001. **j.** Immunostaining for anti-laminin-β1 and anti-GFAP colocalized with GFP^+^ PLCs, in PCCOs. Right is zoom of small region. Bottom shows two-color staining. Bar: 40μm. Single arrow: laminin in gray. Double arrow: astrocytic endfeet connection. **k.** Model for pericytes-basement membrane-astrocytes structure in PCCOs. PLCs express laminin and promote astrocytic maturation and endfeet attachment to PLCs. DAPI (blue) for nuclei.

We found that PCCOs maintained similar structural architecture and cellular compositions as traditional COs (Fig. 2d-f and Extended Data Fig. 2a-b). The GFP^+^ PLCs within PCCOs show *GFP* mRNA expression and retained standard pericyte markers expression (Extended Data Fig. 3a-b). However, we found that within PCCOs, PLCs attuned GFAP-positive astrocyte expression and morphology, which more closely resembled a classically described ‘star’-shape with end-feet like structures seen in mature astrocytes that were adjacent to PLCs (Fig. 2g)^22,23^. These results evidenced features of astrocytic maturation compared to traditional COs^24^. Moreover, we detected laminin-β1 protein adjacent to PLCs, suggesting accumulation of basement membrane (BM), which is normally absent from traditional COs (Fig. 2h-i). Confocal imaging confirmed localization of astrocytes and PLCs with laminin (Fig. 2j). Together, these results suggest PCCOs recapitulate the structural architecture of PLCs-BM-astrocytes described within the vertebral neurovascular unit (Fig. 2k)^16^.

We additionally characterized PCCOs by single cell RNA-seq (sc-RNA-seq). Compared to COs, PCCOs showed an ~23% shift from progenitor to neuronal populations, which was validated with TBR1 immunostaining (Fig. 2e, Extended Data Fig. 4a-d and Supplementary Dataset 1)^25–27^. Tandem mass spectrometry using isobaric labeling of PCCOs compared with COs supported these results, revealing that GFAP, TBR1, DCX, and STMN2 formed an upregulated protein module, suggesting an effect of PLCs on neuronal differentiation in PCCOs (Extended Data Fig. 5a-b, Supplementary Dataset 1).

We next exposed PCCOs to SARS-CoV-2 at MOI 0.5 for 72h (Fig. 3a). Compared with traditional COs that showed scant neuroglial cells positive for the established viral SNP protein as reported^28,29^, PCCOs showed a significantly higher proportion of SNP^+^ cells (10% vs. 1% in PCCOs vs. COs, p <0.0001, t-test, Fig. 3b-f, Extended Data Fig. 6a-b, Supplementary Dataset 2). qRT-PCR showed a corresponding ~50-fold increase in viral RNA in exposed PCCOs over COs (Fig. 3g, Supplementary Dataset 2).

**Fig. 3.**
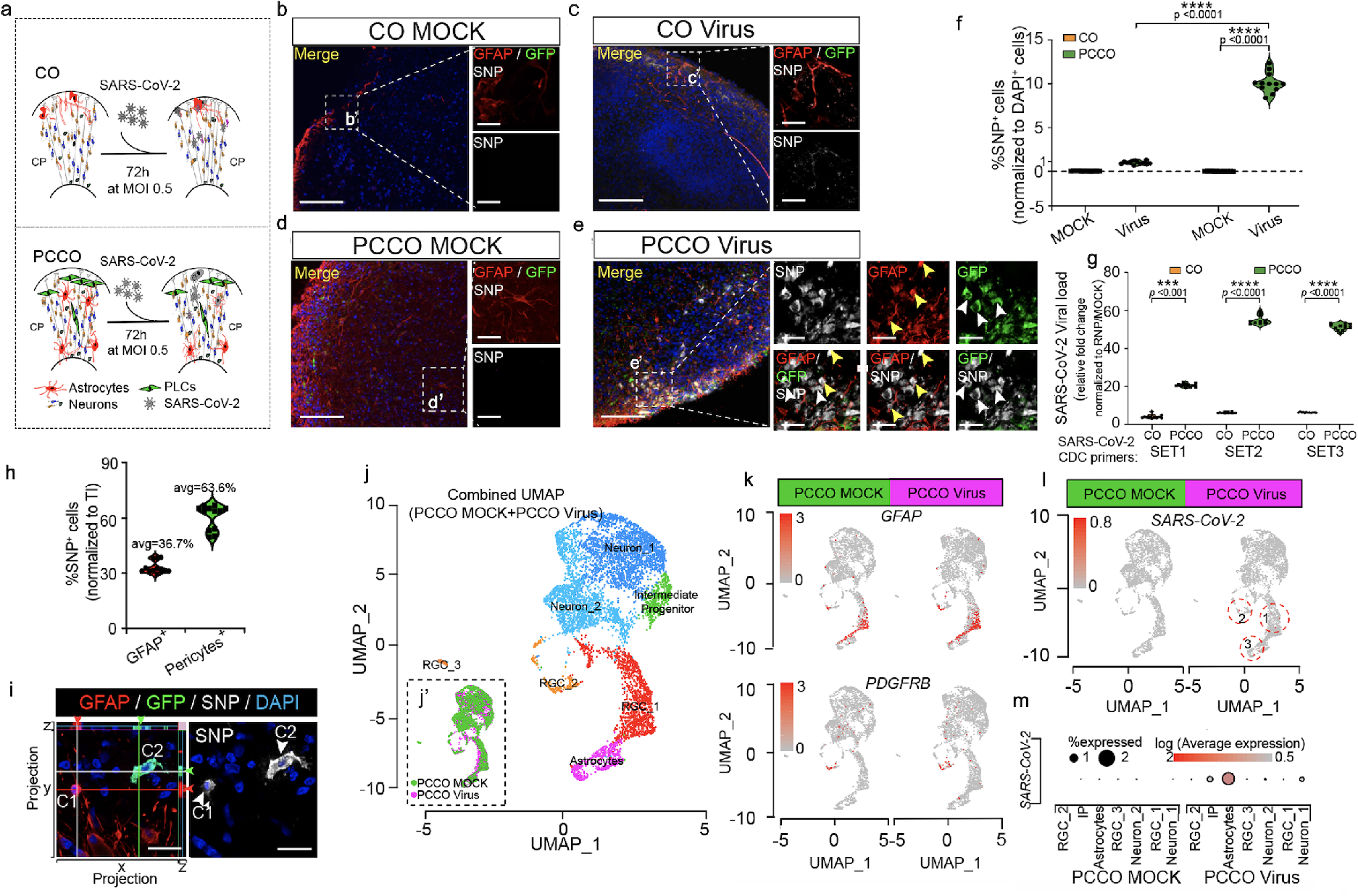
SARS-CoV-2 productively infects PCCOs but not COs. **a.** SARS-CoV-2 infection in COs and PCCOs protocol. **b-e.** GFAP and SNP staining under MOCK and virus infection conditions. GFP: PLCs, DAPI: nuclei, GFAP: astrocytes, SNP: SARS-CoV-2 Nucleocapsid Protein. Dramatic increase in SNP signal seen in PCCOs compared with COs. Bar: 400μm. **b’-e’.** Zoom in panels of B-E. In E’, yellow arrows: GFAP^+^/SNP^+^ double positive i.e. infected astrocyte, White arrows: SNP^+^/GFP^+^ double positive i.e. infected PLCs. Bar: 20μm. **f.** Quantification of SNP positive cells shown in **b-e**, numbers of DAPI (nDAPI) were used as reference. **g.** qRT-PCR using SARS-CoV-2 primers. Increased viral RNA load in PCCOs 72h post infection. MOCK infection used as negative control. **h.** Quantification of SNP positive cells in E’; nDAPI used as reference. All ratios were normalized to total number of infected cells (TI). Approximately 2/3 of infected cells were PLCs and 1/3 were astrocytes. **i.** SARS-CoV-2 (gray) can infect both astrocytes (GFAP^+^, C1) and PLCs (GFP^+^, C2) in PCCOs. Side bars: x-z and y-z max projections demonstrating these are distinct cells. Bar: 20μm. **j-j’.** Merged UMAP shows cellular compositions in PCCOs. 7 cell clusters were annotated according to standard marker gene expression. The split UMAP shows the contribution of PCCO MOCK and PCCO Virus in j’. **k.** Feature plots show the expression of *GFAP* and *PDGFRB* in MOCK and Virus in PCCOs. *GFAP:* astrocytes, *PDGFRB:* PLCs. **l.** Feature plots show *SARS-COV-2* (i.e. any transcript of SARS-CoV-2) expression in PCCOs Viruus. **m.** Dotplots show approximately 2% of astrocytes display evidence of *SARS-CoV-2* infection. In **f**, n=12 includes 3 PCCOs from different biological replicates and 4 image regions from each section. In **g**, n=12 includes 3 biological replicates and 4 technical replicates for each biological replicate. *GAPDH* used as reference. t-test with Sidak multiplecomparison test correction was used to determine the significance. **** p<0.0001.

We then compared the cellular infection vulnerability to SARS-CoV-2 in COs vs. PCCOs. In COs, we found <1% NeuN^+^/SNP^+^ cells or GFAP^+^/SNP^+^ cells and no discernable effect of viral exposure on CO characteristics (Extended Data Fig. 7a-e). In many ways, it seemed that the presence of SARS-CoV-2 was innocuous to COs. In contrast, in PCCOs we found the majority of SNP^+^ cells co-localized with GFP^+^ PLCs and surrounding GFAP^+^ astrocytes in virus-exposed PCCOs (Fig. 3e-e’ and h). Confocal imaging demonstrated that astrocytes were not only adjacent to the infected PLCs but were themselves SNP^+^ (Fig. 3i). To transcriptionally profile cellular constituents, we performed sc-RNA-seq at 72h.p.i. We detected SARS-CoV-2 reads in ~2% of cells in PCCOs, overwhelmingly confined to astrocytes, but not neurons (Fig. 3j-m). There were no detectable SARS-CoV-2 reads in infected COs (Supplementary Dataset 1). These data suggest that infection of astrocytes is mediated by the presence of the PLCs population.

Finally, to explore pathogenesis of SARS-CoV-2 in PCCOs, we performed immunostaining and observed a significant increase (~20%) in percent of cells evidencing programmed cell death (cleaved caspase 3 and p53-positive) in infected PCCOs (Fig. 4a-b, Supplementary Dataset 2). sc-RNA-seq indicated that the source of the cell death was largely confined to astrocytes, consistent with the described selective vulnerability (Fig. 4c)^30^. Gene ontology (GO) term analysis of differentially expressed genes (DEGs) specific to astrocytes highlighted inflammatory and genotoxic stress activation (Fig. 4d). This correlated with activation of type I interferon transcriptional response, and with upregulation of *IFIT1, IFI44, ISG15* in virus exposed compared to mock infected PCCOs (Fig. 4e, Supplementary Dataset 3)^31^. Several type I interferons signaling cascade genes *(STAT1, STAT2*) and an *ISG15-effector* gene *(USP18)* were also upregulated (Fig. 4f, Supplementary Dataset 2)^32^. Increased expression of *ISG15* was confirmed by qRT-PCR in infected PCCOs (Fig. 4g). These results implicate astrocytic pathology in SARS-CoV-2 inflammatory brain pathology, mediated through the type I interferon pathway.

**Fig. 4.**
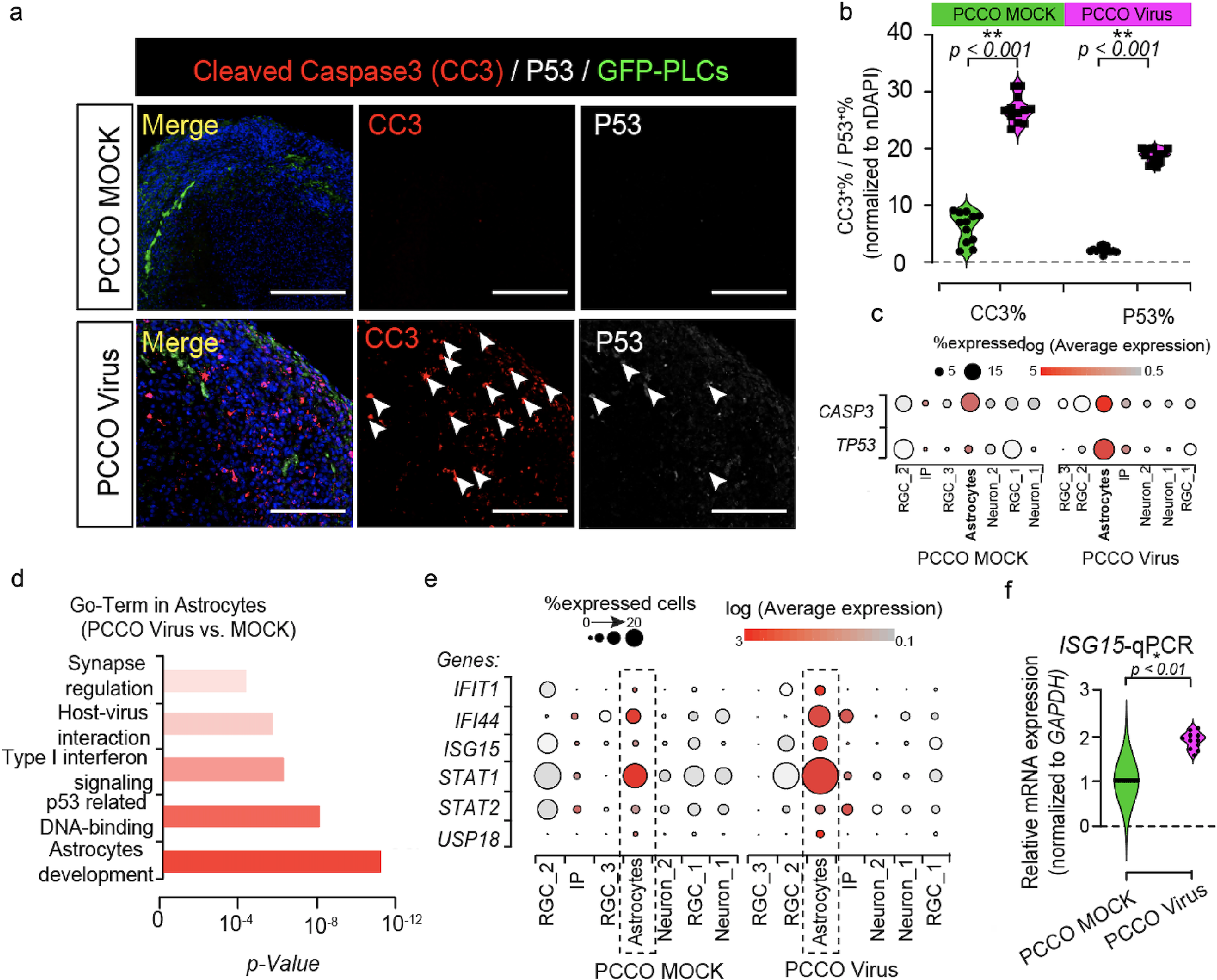
PCCOs SARS-CoV-2 infection elicits type I interferon astrocytic response. **a.** Cleaved Caspase 3 (CC3) and p53 staining is positive in PCCOs following infection. DAPI: nuclei. GFP: PLCs, and P53: gray. Bar: 400μm. Arrow: CC3 or p53 positive cells. **b.** Quantification for a. nDAPI used as reference. **c.** Dot-plots: expression of *CASP3* and *TP53* in PCCO MOCK and Virus groups. **d.** GO-Term with significance for functional analysis in astrocytes. **e.** Dot-plots: expression of type I IFN related genes. **f.** qPCR for *ISG15* mRNA expression; *GAPDH* reference. MOCK infection used as negative control. In **b** and **f**, n=12 includes 3 biological replicates and 4 technical replicates for each biological replicate. *GAPDH* used as reference. t-test with Sidak multiple-comparison test correction was used to determine the significance. ** p<0.001 (**b**), * p<0.01 (**f**). In Dot plots, Dot size: % cells demonstrating gene expression. Dot shade: average expression in log-fold scale.

Here we demonstrate that PLCs can be productively infected by SARS-CoV-2 and through integration of PLCs into COs, we established a novel PCCO ‘assembloid’. Within PCCOs, we found PLCs establish characteristics of PLCs-BM-astrocytes structure and increase the cellular proportion of neuronal population, mimicking reported functions of human brain pericytes *in vivo.* Upon exposure to SARS-CoV-2, we observed robust infection within PCCOs and consequent induction of neuronal death and type I interferon responses. Furthermore, we demonstrate that PLCs can serve as viral ‘replication hubs’, supporting the viral invasion and spread to other cell types, including astrocytes.

Although SARS-CoV-2 invasion into human CNS has been modeled in 3D brain organoids, choroid plexus (ChP) organoids, and K18-hACE2 transgenic mice, evidence suggests that most neural cells have little to no capacity for SARS-CoV-2 infection. On the other hand, the presence of any cells expressing ACE2 or other receptors may be sufficient to initiate infection^28,29,33,34^, motivating further work to understand the receptor expression profile and impact on infection at the human NVU *in vivo.* Drawing on clinical and experimental data supporting potential vascular entry and ACE2 expression in pericytes, our PCCO SARS-CoV-2 infection model presents an alternative route to infection. The PCCO model could be further improved by incorporating other NVU-component cell types, which might lend itself to other uses^35^. Our work provides a powerful model to study SARS-CoV-2, and may be useful to model other infectious diseases.

## Methods

### Human Induced Pluripotent Stem Cells, Neural Crest Stem Cells Culture, and constructs

HEK293T cells (sex typed as female), Hela and H1 ESC (sex typed as male) were obtained from ATCC (CRL-11268™), ATCC (ATCC®CCL-2^TM^) and WiCell (WAe001-A), and were not further authenticated. Generation of neural crest stem cells (NCSCs) was described previously^20^. Human induced pluripotent stem cells (hiPSCs) were from CIRM (CIRM-IT1-06611). All cells were regularly mycoplasma negative. pLV-EF1a-EGFP-Puro construct was obtained from Dr. Fred Gage’s lab as a gift. Lentiviral packaging plasmids pMD2.G, pPAX2 were obtained from Addgene. pcDNA3.1-hACE2 plasmid was obtained from Dr. Tom Rogers at Scripps as a gift.

### Human cortical brain organoid (CO) and pericyte containing cortical organoid (PCCO) culture

H1 and hiPSCs were maintained in mTeSR and passaged according to manufacturer’s recommendations. Cortical brain organoids were generated as previously described^36,37^. For PCCO generation, at CO 60 div, 2×10^5^ GFP+ PLCs were integrated into each CO in a low attachment 96-well plate. PLC integrated COs were maintained in PCCO media for 14 days (See Supplementary Note for details).

### SARS-CoV-2 infection of pericyte-like cells, COs, and PCCOs and Plaque assay

All work with SARS-CoV-2 was conducted in Biosafety Level-3 conditions at the University of California San Diego following the guidelines approved by the Institutional Biosafety Committee. SARS-CoV-2 isolate USA-WA1/2020 (BEI Resources) was propagated and infectious units quantified by plaque assay using Vero E6 (ATCC) cells. See the infection details in Supplementary Note.

### Detection of viral mRNA and replication using RT-qPCR

For viral RNA quantification, PLCs, COs or PCCOs were washed twice with PBS and lysed in TRIzol. RNA was extracted using the Qiagen-RNA extraction Kit. 2μg RNA was used to generate cDNA with SuperScript III First-Strand Synthesis Kit (Invitrogen). 20ng cDNA was used to perform qPCR with iTaq Universal SYBR Green Supermix and the CDC-N1/N2/N3-SARS-CoV-2 primers mix (IDT) at a final concentration of 100nM for each primer using a Bio-Rad Real-Time PCR system. All the qPCR primers were listed in Supplementary Table 1 and Supplementary Datasets.

### Immunostaining and Light Sheet Imaging of PCCO and CO

COs and PCCOs were fixed in 4% paraformaldehyde for 72h before removal from BSL3, then embedded in 15%/15% gelatin/sucrose solution and sectioned at 20 μm. The sections were then permeabilized in 0.5% Triton X-100, blocked with 5% BSA and incubated with primary antibodies in 5% BSA/0.5% Triton X-100 in PBS at standard dilutions overnight at 4 °C. Next day, the sections were incubated with secondary antibodies together with DAPI and mounted with Fluoromount-G®. All the images were taken with ZEISS LSM880 Airyscan, with post-acquisition analysis done in ImageJ-6. For light sheet, PCCOs were harvested for clearing in PBST in a 1.5ml EP tube (1PCCO/tube). CUBIC ^38^ was used for clearing. Cleared PCCOs then embedded into 1% Agarose solution for imaging. 5X lens was used for imaging with a light sheet microscope (Zeiss Z1) according to manufacturer recommendations. The images were further developed with IMARIS (Oxford Instruments).

### CO/PCCO dissociation and Single cell library preparation and sequencing

COs and PCCOs were dissociated using AccuMax, dead cells were removed by Dead Removal cocktail (Annexin V, STEMCELL Technology). Live cells were then used 10X GEM generation. 10X sc-RNA-seq-3’-V3.1 kit (10X Genomics) was used to generate the GEM, cDNA and library were generated according to the manufacturer’s instructions (10X Genomics). Libraries was sequenced using the Novaseq6000 with PE150bp for 20M reads were requested for each sample. See experimental details in Supplementary Note

### CO/PCCO TMT4 quantitative Protein Mass Spectrometry

Three COs and three PCCOs were harvested into 1.5ml cold PBS, then centrifuged at 1500rpm at 4°C for 10min. After centrifugation, all PBS was removed and the CO/PCCO samples were flash frozen in liquid N2. The frozen cell pellets were analyzed by TMT4 quantitative mass spectrometry at the UCSD Proteomics Core.

### Data processing of single-cell RNA-seα and Mass Spectrometry

Single cell RNA-seq sequencing were demultiplexed into Fastq files using the Cell Ranger (10x Genomics, 4.0) mkfastq function. Samples were then aligned to GRChg38-2020 10x genome reference. The count matrix was generated using the count function with default settings. SARS-CoV-2 (USA-WA1/2020) genome and GFP-CDS were written into GRChg38-2020 human genome reference as a gene with mkref function in cell ranger 4.0. Seurat package (v.3.1.5) in RStudio (with R v. 3.5.3), used for the downstream analysis. Features expressed in less than 5 cells, cells with less than 300 unique features, or high mitochondrial content over 5% were discarded. scTransform function was used to wrap the technical variation; FindIntergrationAnchors and IntegrateData functions were used to integrate all libraries metrics into a single matrix. Principal Component Analysis (PCA) together with the PCEIbowPlot function were used to determine the inflection point. Clusters were determined using Runheatmap, FindNeighbors and Findclusters function within Seurat. Cells were considered infected if they carried the transcripts aligned to SARS-CoV-2 viral genome. Differentially expressed genes from the FindMarkers function were used to perform DAVID-GO-Term analysis over representation tests for both up-regulated and downregulated genes in each condition shown in Fig. 2r, Supplementary Dataset 1-2.

Mass Spectrometry data were generated and analyzed by the proteomic core at UCSD (Supplementary Dataset 1). Proteins with significance over 15 were used for GO-Term and string analysis in goprofile (https://biit.cs.ut.ee/gprofiler/gost)and String (https://string-db.org/) (Supplementary Dataset 1).

### Reporting Summary

Further information on research design is available in the Nature Research Reporting Summary linked to this article.

### Data and material availability

The accession number for the sc-RNA-seq data reported in this paper is SRA: PRJNA668200. Further information and requests for resources and reagents should be directed to and will be fulfilled by the Lead Contact, Dr. Joseph Gleeson.

### Quantification and statistical analysis

Statistical analysis was performed with the GraphPad Prism 8 software. We compared viral titer by two-way ANOVA followed by a Sidak multiple-comparison test, * p<0.1. In relative mRNA expression level, cell numbers and all other statistics, n=12 includes 3 from different biological replicates and 4 technical replicates; Multiple t-test was used to determine the significance followed by a Sidak multiple-comparison test correction. **** p<0.00001, *** p<0.0001, **p<0.001, * p<0.01;

## Acknowledgements

We thank Drs. Ronald Ellis, Jeff Esko and Dillon Chen for feedback, Sangmoon Lee for bioinformatics support, UCSD IGM for sequencing support and the UCSD Proteomic Core for mass spectrometry support. The following reagent was deposited by the Centers for Disease Control and Prevention and obtained through BEI Resources, NIAID, NIH: SARS-Related Coronavirus 2, Isolate USA-WA1/2020, NR-52281. The work was supported by the Rady Children’s Hospital Neuroscience Endowment and R01NS106387 to J.G.G, Career Award for Medical Scientists from the Burroughs Wellcome Fund and K08Al130381 to A.F.C., NS103844 to E.V.S. and S.P.P., NIH Biotechnology Training Program T32 GM008349 to B.D.G. National Science Foundation Graduate Research Fellowship Program, 1747503 to B.D.G., and the UCSD Neuroscience Microscopy Core Facility 3P30NS047101 and the UCSD Institute for Genomic Medicine Core Facility 1S10OD026929.

## Author Information

### Contributions

L.W. D.S. H.F. designed and conducted the study. L.W. established the PCCO system using neural crest stem cells provided by B.D.G, S.P.P and E.V.S. A.F.C and A.E.C. performed virus infection, plaque assay and generated single cell RNAseq GEM libraries; L.W., D.S., A.F.C and J.G.G wrote the manuscript; A.F.C and J.G.G supervised the project. All authors reviewed the manuscript.

